# Time-frequency EEG markers of word boundaries in speech production

**DOI:** 10.64898/2026.03.23.713128

**Authors:** Sophie Drew Eustace, Sara Guediche, Lorenza Brasiello, Mónica Rocha, João Mendonça Correia

## Abstract

Speech production requires orchestration of multiple brain systems, including cortical and subcortical areas that support the unfolding of the spoken message across hierarchical linguistic levels, such as phonemes, syllables, words or phrases. Transitions between levels are critical for fluent speech, yet the neural dynamics of, for example, syllable-level and word-level transitions remain unknown. In this electroencephalography (EEG) study, we use time-frequency analysis and source localization to determine differences associated with word-boundary vs. within-word syllable transitions. To this end, pseudoword pairs comprising six consonant-vowel (CV) syllables with different word-boundary positions were used. Fluent human adults produced the utterances at the rhythm of a learned visual metronome (i.e., syllable-by-syllable), such that each syllable was uttered at matching times independently of its relative word position. Accordingly, a target syllable could be either a within-word syllabic transition or a between-word transition, while other linguistic properties, including articulation, stress pattern, co-articulation or prosody, were matched. EEG time-frequency analyses of neural sources successfully revealed sensitivity to hierarchical structure. Neural sources in left and right inferior frontal lobes, as well as left superior temporal lobe were differentially recruited when producing the same exact syllables, in the same exact utterance position, but under different word boundary contexts. A right inferior frontal source showed a robust time-frequency modulation in word transitions that included elevated event-related synchronization in the theta and beta range. Interestingly, despite our efforts to control speech pace across conditions using metronome-based guidance, small, albeit significant timing delays emerged, confirming higher cognitive demands at word boundaries.

## 1. Introduction

Language is an organised communication system in which smaller linguistic units are embedded within larger structures: phonemes form syllables, which combine to form words, and words can be used to build phrases and sentences (Rauschecker and Scott, 2009; Peelle and Davis, 2012). This hierarchical structure is reflected in the neurobiology of both speech perception (Ding *et al*., 2015) and production (Sengupta and Nasir, 2016; Tremblay, Deschamps and Gracco, 2016; de Heer *et al*., 2017) and supported by temporally distinct patterns of neural activity (Giraud and Poeppel, 2012; Peelle and Davis, 2012). A speech perception study (Ding *et al*., 2015) used isochronous streams of syllables that could be grouped into words, phrases or sentences at different fixed rates, and showed that cortical electrophysiological activity synchronised (i.e., entrained) not only to the underlying syllabic rhythm but also to the word, phrase and sentence rhythms. This frequency-tagging paradigm demonstrates that neural oscillations within posterior temporal and inferior frontal cortices track the hierarchical linguistic structure of speech perception. In other words, the brain may have special neural signatures at different levels of speech hierarchy. Similarly, continuous speech production involves transforming abstract linguistic representations into precisely timed articulatory actions, engaging cortical and subcortical regions. Foundational neurocognitive models, such as the ‘Blueprint of the Speaker’ (Indefrey and Levelt, 2004) or the dual-stream framework (Hickok and Poeppel, 2007; Hickok, 2012) also describe speech production using a similar hierarchy. Overall, brain regions including motor and premotor cortices are associated with motor planning and execution, auditory cortices with sensory-feedback prediction and monitoring of speech outcomes (Houde and Jordan, 1998; Tian and Poeppel, 2010; Christoffels *et al*., 2011), and basal ganglia and cerebellar structures with timing and sequencing of motor programmes (Knolle *et al*., 2019; Archila-Meléndez *et al*., 2020; Jorge *et al*., 2022). These regions may play different roles at different hierarchical levels of speech.

In motor-speech control, an efference copy (i.e., a forward model) is an internal representation of the sensory consequences of a planned motor command (Crapse and Sommer, 2008), which enables the rapid evaluation of incoming sensory feedback for error detection and correction (Ford, Roach and Mathalon, 2010; Niziolek, Nagarajan and Houde, 2013; Simmonds *et al*., 2014). At present, a key challenge of speech production is disentangling different levels of planning from production proper and monitoring of predicted sensory consequences (Tremblay, Deschamps and Gracco, 2016). Producing any utterance involves a continuously updated planning scope that spans upcoming segments and often the next word. Consequently, neural activity around a speech-production boundary (e.g., a syllable or word) may reflect (i) syllabic sequencing and articulatory chunking, (ii) lexical selection and word-level planning, and (iii) sensory predictions and error monitoring, all partially overlapping in time. Hence, to study the full-scale neural processes involved in speech production requires continuous speaking paradigms, while experimentally isolating the different neural processes involved. Interestingly, debate remains on what are the basic units of articulatory planning. Some studies suggest that syllables are the units of articulatory planning (Laganaro, Valente and Perret, 2012; Bürki *et al*., 2016), whereas others postulate that speech planning can proceed at the word level (Piai *et al*., 2015; Assaneo and Poeppel, 2018).

Distinguishing between speech planning, monitoring and execution processes may also shed light on the neural disruptions present in motor-speech disorders, including stuttering, where fluent transitions between speech units are compromised (Indefrey, 2011; Chang and Guenther, 2020). Interestingly, the relative position of speech gestures within words affects stuttering behaviour. Studies have shown considerably higher dysfluency rates at word, phrase and sentence boundary transitions, compared to within-word syllabic transitions (Bloodstein and Ratner, 2008; Buhr and Zebrowski, 2009). This dysfluency disproportion could arise from either encoding, prediction or monitoring (Loucks *et al*., 2007; Ghazanfar and Eliades, 2014). Furthermore, the basal ganglia play a critical role in motor control by modulating the cortex for motor initiation/execution, based on the direct and indirect striatum-based pathways (Schulz *et al*., 2005). These cortico-basal ganglia-thalamo-cortical circuits (aka, the cortico-basal ganglia loop) are essential for the execution of motor commands, including motor speech gestures. Their role in stuttering has been suggested, as well as, possible parallels between stuttering and Parkinson’s disease (PD) (Alm, 2004; Giraud *et al*., 2008). Beyond the possible role of the direct and indirect pathways in stuttering, a more recent link between stuttering and the basal ganglia has been proposed based on failures within a faster cortico-basal ganglia loop that short-cuts the striatum, and instead stimulates the sub-thalamic nucleus directly - the hyperdirect pathway (HDP) (Neef, Anwander and Friederici, 2015). The HDP leads to hyper-fast inhibitory signals to the cortex (Nambu, Tokuno and Takada, 2002). Precision in inhibitory motor control via the HDP is suggested as necessary to stop or correct ongoing motor actions, thereby contributing to sensorimotor control in speech production (Whillier *et al*., 2018; Usler, 2022). Importantly, the HDP may also be critically involved in motor/speech initiation (Usler, 2022; Nambu *et al*., 2023) by resetting the primary motor cortex (M1) prior to the arrival of the excitatory direct pathway that ultimately leads to muscle contraction (Nambu, 2011). In turn, cortical inhibition by the HDP may be particularly important at higher hierarchical levels of motor planning requiring M1 reset, including transitions between words, phrases and sentences. Importantly, impairments along the HDP has been implicated in stuttering (Usler, 2022).

Transitions between linguistic units, whether within a word (e.g., syllable-to-syllable) or between words (e.g., word-to-word), are hypothesized to engage distinct neural oscillatory mechanisms (Cao, Thut and Gross, 2017; Cao *et al*., 2024; Orpella *et al*., 2024). Prior studies have used delayed-production paradigms (Kittilstved *et al*., 2018; Archila-Meléndez *et al*., 2020), in which speech planning and production proper (i.e., speech execution) are separated by an experimental cue. While this approach helps to dissociate planning- from execution-related activity, it limits our understanding of how neural dynamics support online transitions during continuous speech production. In addition, while it is possible to instruct participants to produce isolated speech units of different hierarchical levels (e.g., isolated syllables versus words) (Silbert *et al*., 2014), probing these mechanisms in sequential/continuous speaking tasks is crucial. For instance, producing the syllable *<u>ba</u>* in different contexts: i) in isolation; ii) as the first syllable in the word *<u>ba</u>nana*; or iii) as the final syllable in the word *scu<u>ba,</u>* may involve distinct neural processes (from planning to monitoring) for the *ba* component, despite requiring the same broad articulatory gestures. However, these conditions hold co-articulatory differences that confound how the syllable *<u>ba,</u>* in this example, is produced in the sequence context. If, instead, target syllables are embedded in a syllabic sequence with matching co-articulatory conditions, but under different word-boundaries, it becomes possible to study hierarchical speech production processes in a controlled manner. We attempt to provide an analogy of how we to test this question using English language: if the word *‘scu’ and ‘nana’* would be legal english words, the sequences ‘*scu<u>ba</u> nana*’ and ‘*scu <u>ba</u>nana*’ would be valid phrases allowing to study how the target syllable *<u>ba</u>* plays different hierarchical roles in the sequences, while preserving its phonological/phonetic form and co -articulatory effects. In practice, working with pseudoword pairs enables the construction of controlled stimulus sequences (that occur across different syllable positions) across experimental conditions, which we use in this study.

EEG time-frequency analyses, such as event-related spectral perturbations (ERSP) allow tracking transient power changes across frequency bands in response to time-locked experimental events (Delorme and Makeig, 2004), which can be used to separate hierarchical transitions and explore the neural dynamics of speech production (Vos *et al*., 2010; Jenson *et al*., 2014). Beta oscillatory activity (∼15-30 Hz) over frontal cortices - often prominent over right inferior frontal and premotor regions - has been associated with preparatory set/maintenance and sequential control during speech and orofacial actions (Pfurtscheller and Lopes da Silva, 1999; Weiss and Mueller, 2012). Converging evidence indicates a functional subdivision whereby low beta (∼15-20 Hz) is more closely associated with the indirect basal ganglia pathway whereas high beta (∼20-30 Hz) reflects hyperdirect cortico-subthalamic signals, coupling the premotor cortex (e.g., inferior frontal regions - IFG, specially the right IFG) and the supplementary motor areas (SMA) with the STN (Oswal *et al*., 2021; Herz *et al*., 2023; Cao *et al*., 2024). Changes in time-frequency provide neural signatures for different aspects of speech production, which may help unveil the neurobiological failures that contribute to stuttering, mainly around word-boundaries. Clarifying how oscillatory signals encode word-to-word transitions in continuous speech may therefore open new venues to understand different neural dysfunctions that can contribute to future research on motor-speech disorders (Neef *et al*., 2018).

In the present study we investigate EEG time-frequency changes during paced overt speech production to compare within-word and between-word syllabic transitions, in the context of pseudowords. Fluent participants produced pseudo-word pairs forming a sequence of six consonant-vowel (CV) syllables, at the pace of a visual metronome. Importantly, pseudo-word pairs consisted of either a two-syllable word followed by a four-syllable word (i.e., condition 2+4) or two three-syllable words (i.e., condition 3+3). This experimental design enabled the direct comparison of neural activity at syllable-to-syllable and word-to-word transitions in a controlled and time-locked manner. In turn, participants produced the same exact syllables, in the same exact sequence position, but under different relative positions within a word ( i.e., first versus last syllable). Accordingly, we focus on EEG time-frequency changes time-locked to the production of the third syllable, as this syllable is an event that corresponds to a word-to-word transition in the 2+4 condition and a within-word syllable-to-syllable transition in 3+3 condition.

## 2. Methodology

### 2.1 Participants

Twenty-one fluent Portuguese-speaking adults (14 females, 5 males), aged between 18 and 53 years (Mean ± Standard Deviation (SD) = 29.4 ± 9.6), participated in the study. Two participants were excluded due to technical errors during the EEG-audio recordings. All participants were right-handed, had completed secondary education or higher education (bachelor’s, master’s, or doctoral level) and reported normal hearing and no history of psychiatric, neurological, or language -related disorders. Participation was voluntary, and no compensation was given for taking part in the study. The study was approved by the ethical committee of University of Algarve, Portugal, and performed in accordance with the Declaration of Helsinki and Oviedo Convention. All participants provided written informed consent prior to testing.

### 2.2 Stimuli

Stimuli consisted of sequences of two pseudo-words designed to isolate neural activity related to syllable- and word-level transitions (see Figure 1B). Each sequence contained two pseudo -words and conformed to one of two structures: condition 2+4 (e.g., *náfa dacalána*), in which a disyllabic pseudo-word was followed by a tetrasyllabic pseudo-word, and condition 3+3 (e.g., *náfada calána*), in which two trisyllabic pseudo-words were presented. Both structures were matched for length (six syllables per trial). All syllables followed a CV structure (‘ba’, ‘da’, ‘fa’, ‘ma’, ‘na’, ‘sa’, ‘ta’, ‘ca’, ‘la’, and ‘ga’) corresponding to the International Phonetic Alphabet (IPA) consonant sounds /b/, /d/, /f/, /m/, /n/, /s/, /t/, /k/,/l/, and /g/ followed by the vowel /a/. We minimised bilabial closures at and before the target syllable 3 to reduce perioral EMG while maintaining clear tongue-tip versus tongue-dorsum articulations. Labial consonants were restricted to speech positions after syllable 3; i.e., no labial articulations occurred at syllable position 2 or 3 to reduce EMG artifacts on the EEG signal (Whitham et al., 2007; Stepp, 2012). The third syllable was always tongue-driven (using /d/ or /k/ consonants). The pseudo-words were phonotactically legal in Portuguese and carried stress on the first syllable of the first pseudo-word and the penultimate syllable of the second pseudo -word, mirroring natural stress patterns while avoiding lexical or semantic familiarity effects (Arruda et al., 2023). Hence, the stress pattern of the individual syllables was the same for all stimuli (and across conditions). Finally, written stimuli were presented in lowercase text using a uniform sans -serif font (size 24), centred on the screen with a grey background using Presentation® software (Neurobehavioral Systems, Inc., Berkeley, CA, USA; www.neurobs.com).

**Figure 1.**
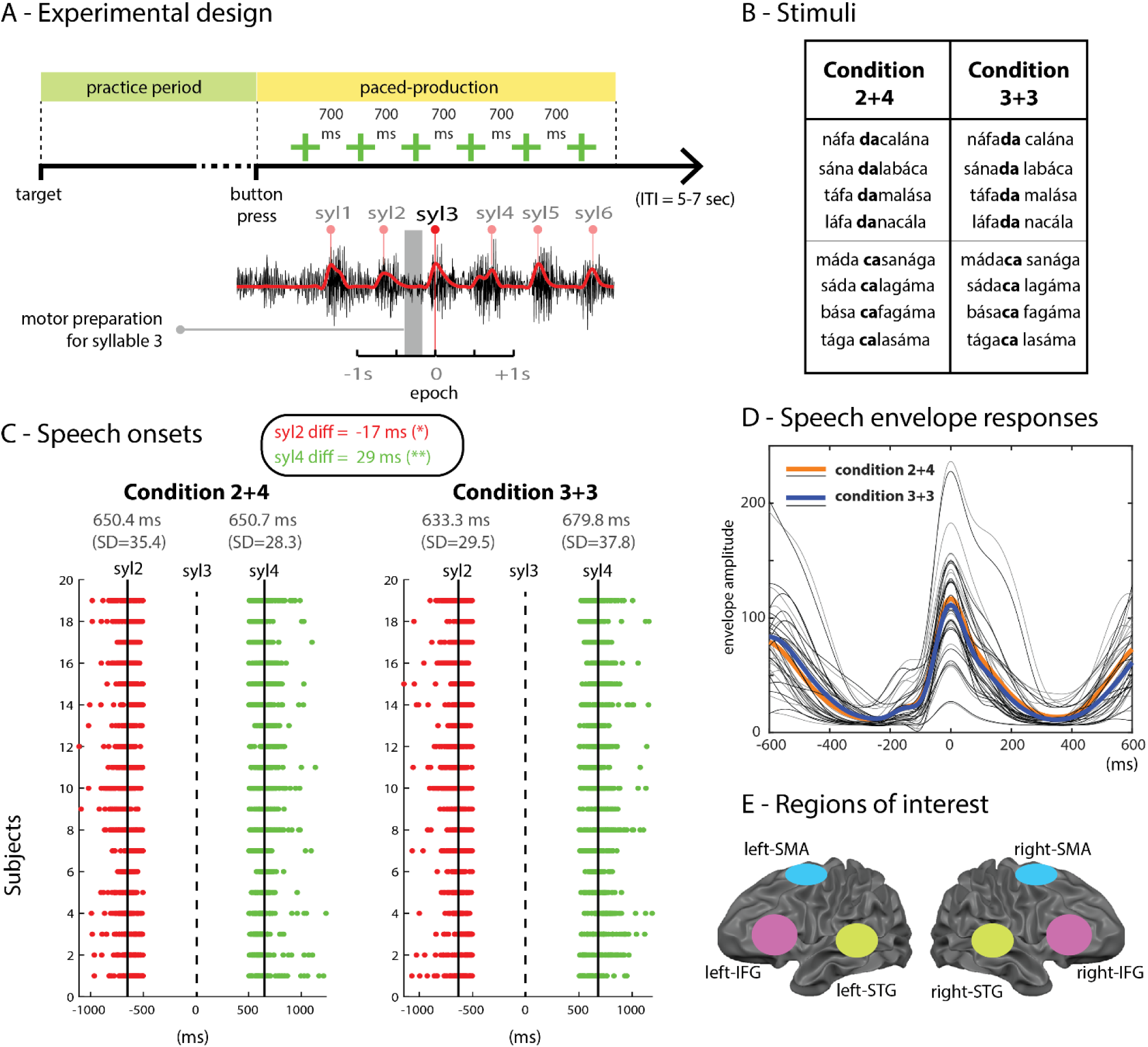
A) Experimental design. ITI – inter-trial-interval. B) Stimuli. Two sequence types were used: Condition 2+4 and Condition 3+3. C) Behavioural results corresponding to speech onsets flanking syllable 3. Red and green dots depict individual trial syllable speech timings, relative to syllable 3, per subject (red = syllable 2; green = syllable 4); vertical lines mark the metronome timings. Mean (SD) speech timings are shown at the top. D) Averaged speech-envelope responses. Group speech envelope traces time-locked to syllable 3 (grey = individual participants; coloured = condition averages). E) Regions of interest. Bilateral inferior frontal gyrus (IFG), superior temporal gyrus (STG), and supplementary motor areas (SMA) used for hypothesis-driven visualisation in ICA cluster selection.

### 2.3 Experimental Procedure

The experiment took place in an electrically shielded EEG laboratory at the Cognitive Neuroscience Group of University of Algarve. EEG data were recorded using a 128-channel BioSemi ActiveTwo system (BioSemi B.V., Amsterdam, the Netherlands) with electrodes positioned according to the international 10/20 system. The vertex sensor (i.e., Cz) was centred between the inion and the nasion anatomical fiducials on the sagittal plane, and the midpoint between the ears on the coronal plane. Conductive gel was applied to each electrode site using a blunt-tipped plastic syringe, and electrode impedance was verified to minimize offsets, in line with BioSemi specifications. Once EEG preparation was completed, participants seated comfortably in front of a computer screen at approximately 70 cm viewing distance.

The experiment consisted of 160 trials in total, divided into 4 blocks of 40 trials each. Each block contained five repetitions of 4 sequences from condition 2+4 and 4 of condition 3+3, fully randomised. Within a block, the same sequence was never presented under both conditions. Hence, blocks differed in how syllables were grouped, providing variation while preserving the same underlying syllabic material across blocks. The order of blocks was counterbalanced across participants, ensuring that all participants completed both conditions under comparable exposure and repetition constraints. Following EEG cap placement and impedance checks, participants began each trial with a practice phase followed by a paced production using a visual metronome (see Figure 1A). The practice phase had the target pseudo -word pair written on the screen, and participants repeatedly uttered the target sequence three times. Upon completion of the trial practice phase, participants pressed the ‘space bar’ button to signal their readiness to proceed to the paced production phase. In the paced production phase, the written sequence disappeared from the screen, and a green fixation cross appeared, flashing every 700 ms (1.43 Hz). Participants were instructed to produce one syllable per flash, aligning their speech with the visual rhythm. Because the lexical boundary shifted across sequence structures, the transitions flanking syllable 3 differed by condition. In condition 2+4, the transitions between syllables 2 and 3 were between-word transitions and in condition 3+3, were within-word transitions. Conversely, the transitions between syllables 3 and 4 were within-word transitions in condition 2+4 and between-word transitions in condition 3+3. This counter balancing enabled within-item contrast of between-word vs within-word syllabic transitions while controlling segmental content and pacing .

### 2.4 Data Recording and Pre-processing

Speech responses were simultaneously recorded with EEG at 16384 Hz (16-bit resolution) using a Shure SM58 microphone placed approximately 15 cm from the participants’ mouth. Continuous EEG was recorded from the 128-channel BioSemi ActiveTwo system (BioSemi B.V., Amsterdam, the Netherlands), with a sampling rate of 512 Hz. Electrodes were arranged according to the international 10/20 system. The BioSemi CMS/DRL system served as the reference and ground during acquisition. Audio data were processed to identify speech response onsets. First, speech recordings were subjected to a Hilbert transformation, and the resulting amplitude envelope was low-passed filtered at 8 Hz and resampled to 256 Hz. Within each trial, six local envelopes (one per syllable) were detected to provide syllable-level temporal markers for identifying the production of the 6 syllables composing our stimuli. Custom Matlab routines were used for audio processing. EEG preprocessing was performed in EEGLAB (Delorme and Makeig, 2004), including custom-made MATLAB scripts (version R2021b; MathWorks, Natick, MA). The continuous EEG recordings were i) resampled to 256 Hz and ii) high-pass filtered at 1 Hz to remove slow signal drifts. iii) Noisy and flat-line channels were identified and removed using the ‘clean_rawdata’ EEGLAB function, after which data were iv) re-referenced to the average of all remaining channels. Across datasets, a mean of 97.9 channels (*SD* = 9.6) remained following bad-channel rejection. Next, v) independent component analysis (ICA; INFOMAX) was performed, and components reflecting ocular, muscular, or other stereotypical artefacts were automatically identified and removed using ‘ICLabel’ and ‘ICFlag’ EEGLAB functions (Pion-Tonachini, Kreutz-Delgado and Makeig, 2019). vi) Channels previously excluded were subsequently interpolated back into the dataset using the ’interp’ method. vii) Following ICA cleaning, the data were low-pass filtered at 70 Hz and a band-stop (48-52 Hz) filter was applied to further suppress line-power noise (i.e., 50 Hz energy). Next, we segmented the continuous EEG into epochs from -1 to +1 secs centred on the third-syllable production event. Time-locking our EEG analysis to the third syllable minimises contamination from sequence-onset preparatory motor activity, while still providing symmetric context for preceding and following syllable transitions. Although the speech envelope peak can lag the true acoustic onset by a small amount, the same alignment procedure and fixed pacing were used in both conditions, and behavioural timing (Figure 1D) confirmed consistent alignment across participants. Event-related spectral perturbations (ERSPs) were computed for each trial and independent component (IC), providing a time-frequency representation of oscillatory time-frequency changes across conditions.

### 2.5 Data Analysis and Statistics

#### Independent Component Analysis and Source Clustering

Equivalent current dipoles were estimated for each independent component using a standard boundary element head model. Components with residual variance greater than 15% were excluded. Next, dipoles were computed for each IC component and participant. Finally, 20 clusters were made by matching dipole locations and spectral activity between 3 and 40 Hz (see Figure 2). Cluster sizes ranged from 19 to 197 ICs (M = 93.1, SD = 40.0). With this approach, we took advantage of the high-density EEG montage (i.e., 128 channels), by performing group statistics at the EEG source level instead of relying on EEG-channel signals, which are known to have low spatial specificity (Delorme & Makeig, 2004).

**Figure 2.**
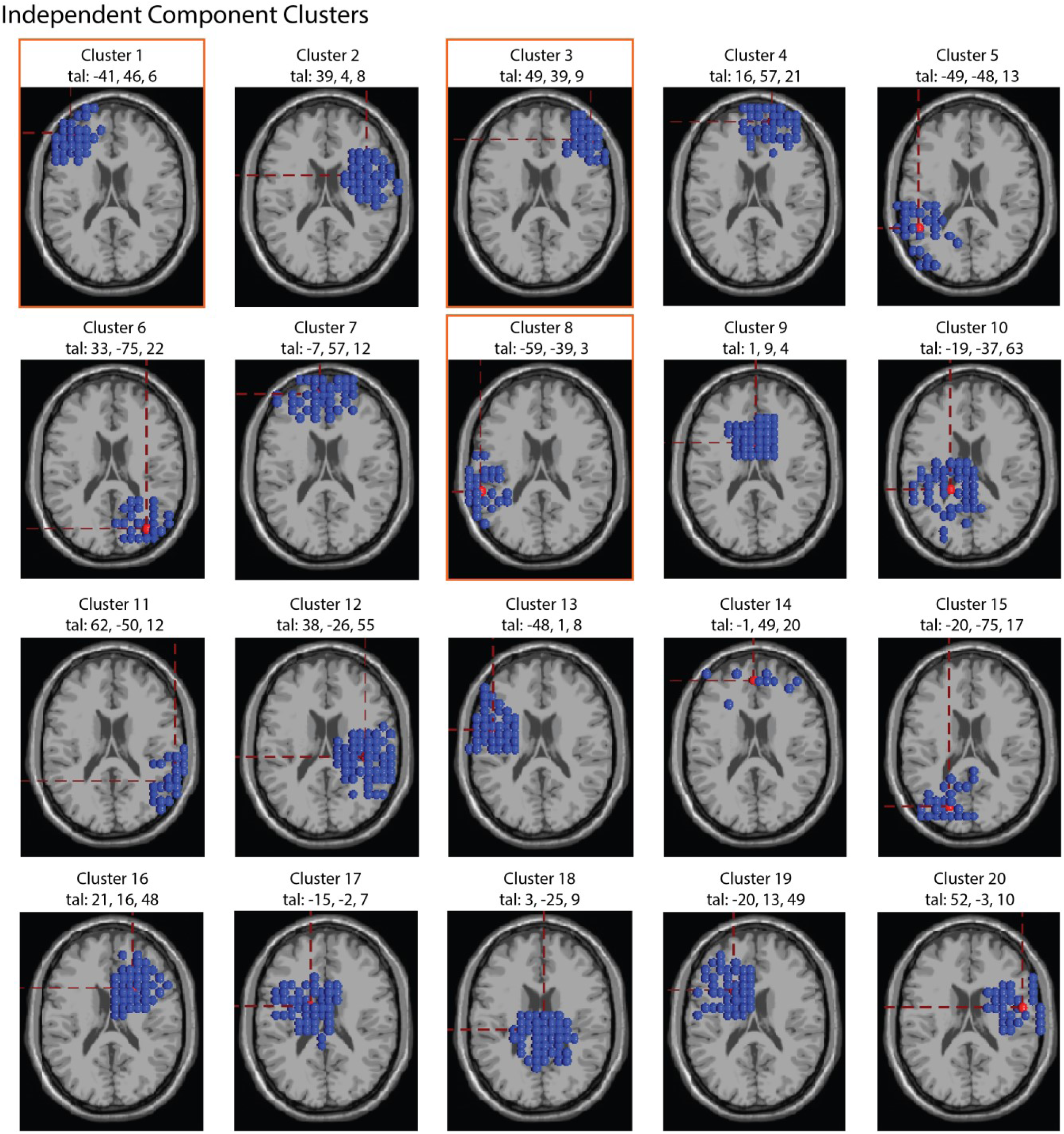
Independent Component (IC) clusters, and centroid location (red sphere) in Talairach coordinates. Blue spheres are individual ICs forming each cluster.

#### Time-Frequency Analysis

EEG was segmented into epochs from -1 to +1 secs aligned to the speech response time of the third syllable (target syllable) of the sequence. Time-locking our EEG analysis to the third syllable minimises contamination from sequence-onset preparatory motor activity, while still providing symmetric context for preceding and following syllable transitions. Although the speech -envelope peaks may lag true acoustic onset by a small amount, the same alignment procedure and fixed pacing were used in both conditions, and behavioural timing (Figure 1D) confirmed consistent alignment across participants. Event-related spectral perturbations (ERSPs) were computed for each trial and independent component (IC), providing a time-frequency representation of time-frequency power changes across conditions.

ERSPs were averaged within each cluster and assessed statistically using a permutation test (based on 2,000 condition-label permutations). For hypothesis-driven visualisation, we report clusters corresponding to bilateral IFG, bilateral STG, and bilateral SMA (Figure 1E); example ERSPs are shown for L-IFG (Cluster 1), R-IFG (Cluster 3), and L-STG (Cluster 8) (see figure 3). ERSPs for all 20 clusters are provided in figure 5. Multiple comparisons over the time-frequency bins were corrected using FDR (*q* = 0.05) (Benjamini and Hochberg, 1995) (i.e., 200 time-points by 68 frequencies). Behavioural contrasts were evaluated with two-tailed, paired t-tests. We refer to decreases in band-limited power as event-related desynchronisation (ERD) and increases as event-related synchronisation (ERS), reporting frequency bands and time windows relative to the epoch average (Delorme and Makeig, 2004).

**Figure 3.**
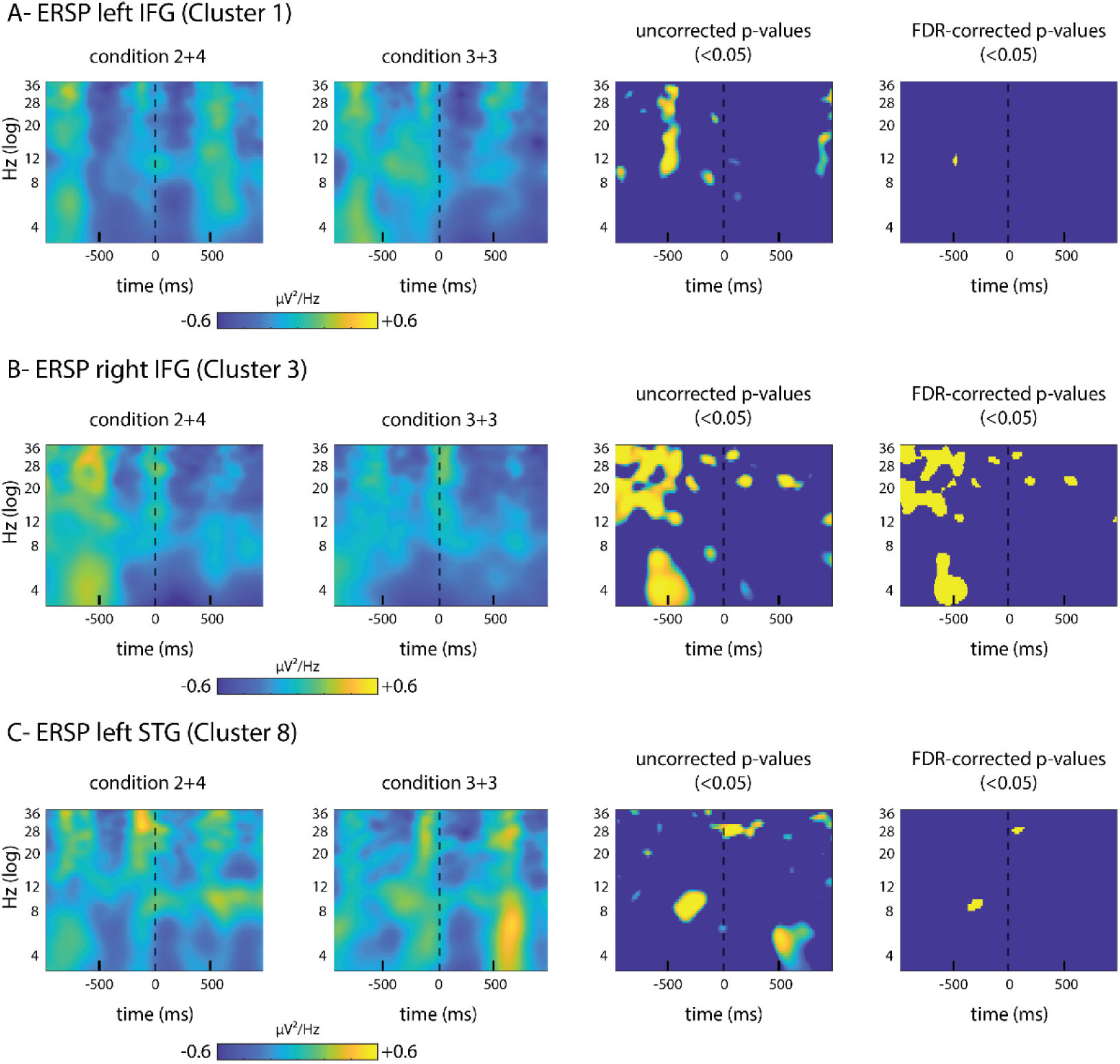
ERSPs in the three clusters matching the selected ROIs (Clusters 1, 3, and 8). First and second left time-frequency plots depict the grand-averaged ERSPs per condition. Third and fourth plots depict ERSP statistics: uncorrected effects (p<0.05) and FDR-corrected (*q* = .05). A) left-IFG cluster. B) right-IFG cluster. C) left-STG cluster.

## 3 Results

### 3.1 Behavioural Results

Participants successfully aligned their speech production with the visual metronome (see Figure 1C-D). Timing between successive syllables around syllable 3 differed slightly, but significantly by condition. The interval between syllable 2 and 3 was significantly shorter in the 3+3 condition in comparison to the 2+4 condition, mean (SD) = 17 ms (34.49), *t*(18) = 2.15, *p* = .045. The interval between syllable 3 and 4 was longer in 3+3 than in 2+4, mean (SD) = 29 ms (43.75), *t*(18) = -2.90, *p* = 0.010. Group speech-envelope traces time-locked to syllable 3 confirmed consistent alignment of the third-syllable envelope peak across participants and conditions (Figure 1D).

### 3.2 Time-frequency results (Event-Related Spectral Perturbations)

Time-frequency differences were analysed within the three ROI clusters (Clusters 1, 3, and 8; see Figure 2). EEG statistics are based on permutation-based condition tests (2,000 label permutations) with FDR for multiple comparisons correction, across time-frequency bins (*q* = .05). Consistent with our analysis plan, bilateral IFG, bilateral STG, and bilateral SMA were treated as confirmatory ROIs (FDR correction within time-frequency bins). No other ROI clusters yielded FDR-corrected effects. For completeness, ERSPs for all 20 clusters are shown in figure 5.

The left IFG (Cluster 1; tal: -41, 46, 6) showed largely similar ERSPs across conditions, with only a small, isolated FDR-corrected significant alpha patch (∼ 10-14 Hz) about 450 ms prior to the target syllable 3 (Figure 3A). Broader uncorrected differences (p<0.05) did not survive FDR correction. In the right IFG (Cluster 3; tal: 49, 39, 9), a robust pre-pk3 modulation survived FDR correction, spanning ∼ 4-30 Hz (theta/alpha into low beta; Figure 3B, FDR panel). Power modulations were greater for the 2+4 condition in comparison to the 3+3 condition, with additional weaker patches evident at the uncorrected level in the early post-syllable-3 window. In the left STG (Cluster 8; tal: -59, -39, 3), we observed two brief FDR-significant time-frequency patches (figure 3C): a pre-syllable 3 alpha patch (∼ 9-11 Hz, [-400 to -300] msecs) and a short post-syllable 3 high-beta patch (∼ 26-30 Hz, [0 to +100] msecs). Outside these windows, no sustained corrected effects were present. The right STG showed no FDR-corrected differences (see figure 3 and 4). Considering all 20 clusters, the clearest FDR-corrected modulation emerged in right IFG, indicating a right-lateralised inferior frontal preparatory signature preceding the target syllable 3. Left IFG and left STG contributed only limited, and no other clusters yielded FDR-corrected effects. See figure 4 for a summary of the significant ERSP contrasts in the confirmatory ROIs.

**Figure 4.**
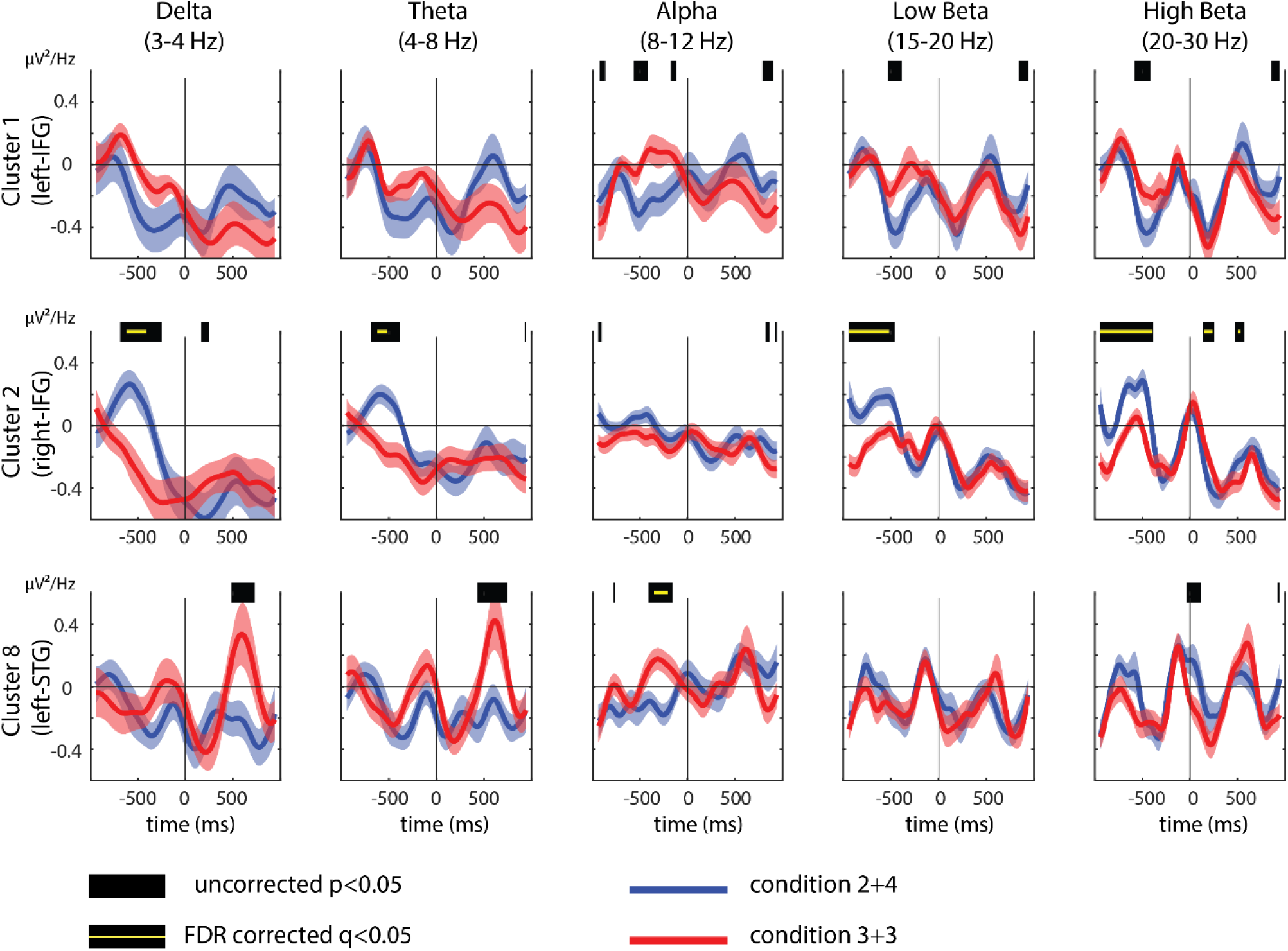
ERSP summary per frequency band (delta, theta, alpha, low beta and high beta) per cluster (cluster 1 – left IFG, cluster 2 – right IFG, cluster 8 – left STG). Blue line is condition 2+4 and red line is condition 3+3. Shaded areas around the blue and red lines represent standard errors from the group mean. Black horizontal bars on top of each cluster-band plot depict uncorrected p values (p<0.05), and the yellow line depict FDR corrected p values (q<0.05).

**Figure 5.**
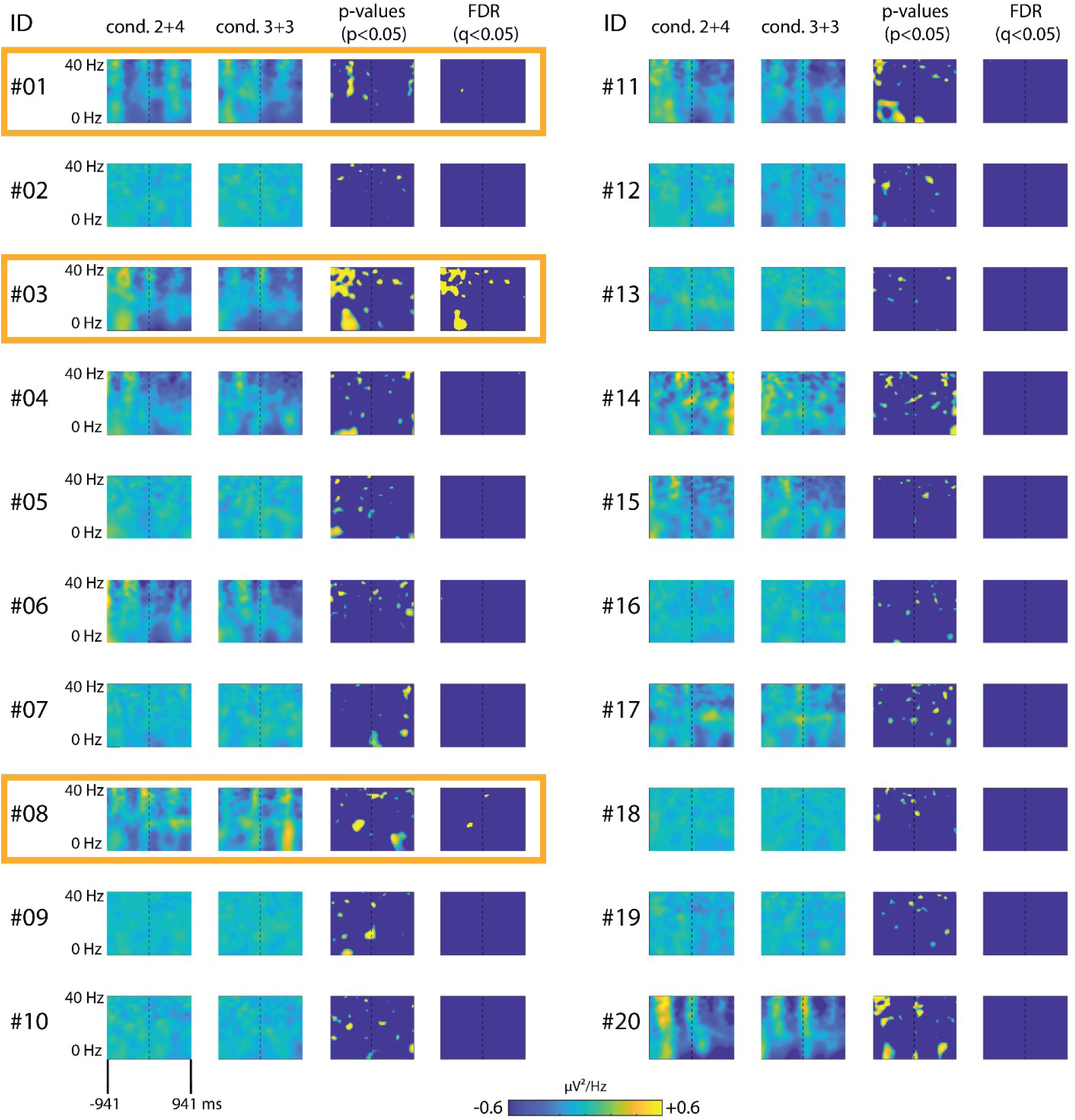
ERSPs in all the 20 clusters (see Figure 2 for cluster locations). First and second left time-frequency plots depict the grand-averaged ERSPs per condition. Third and fourth plots depict ERSP statistics based on permutation testing (2,000 label permutations): third plot is uncorrected effects (p<0.05), and fourth plot is FDR-corrected (*q* = .05).

## 4 Discussion

Although language is a hierarchically organized communication system, segmental boundaries in natural speech are often concealed due to coarticulation. The perceptual challenge for a listener is to recover speech boundaries from continuously presented speech , which involves specific neural oscillations (Ding *et al*., 2015). In speech production, the same hierarchical levels become relevant for understanding motor speech disorders such stuttering, but their neural signatures remain underspecified (Neef *et al*., 2015). Here, we investigated differences in cortical oscillations across different levels of hierarchy-syllable forming word boundary vs. within-word syllable boundary. Clustering of ICA components, obtained from high-density EEG, identified three clusters within our regions of interest, which showed significantly different time-frequency signatures when comparing within-word and between-word syllabic transitions: the left and right IFG and the left STG (Figure 2). The importance of these clusters has been previously identified in the literature, as a distinguishing part of the language and speech network that supports speech planning versus monitoring, respectively (Hickok, 2012; Tremblay, Deschamps and Gracco, 2016). Across these clusters, the most robust differences in modulation were in the right IFG spanning both theta and beta oscillatory bands (Figure 3).

We observed a robust effect spanning theta and beta oscillations around 500 ms prior to the target syllable, which was a between-word transition in the 2+4 condition and a within-word syllabic transition in the 3+3 condition. This effect indicates that frontal control networks adjust preparatory states according to the hierarchical role of the upcoming syllable, maintaining within-word chunking versus reconfiguration at a boundary. This pattern aligns with reports that right-frontal beta oscillations index preparatory and sequential control during speech (Pfurtscheller and Lopes da Silva, 1999; Weiss and Mueller, 2012; Piai *et al*., 2015). Furthermore, theta oscillations may contribute to joint operations with beta rhythms, namely in sensory prediction (Arnal and Giraud, 2012) or cortico-basal ganglia circuits (Oswal *et al*., 2021). Importantly, the right inferior frontal cortex is central in inhibitory control via hyperdirect projections to the subthalamic nucleus, which is expressed in beta neural rhythms (Nambu, 2011; Jorge *et al*., 2022).

These findings have important implications for future research on motor speech disorders, particularly stuttering (Rocha, Carmona and Correia, 2025). In stuttering, sensorimotor processes that support fluent transitions between speech units are often disrupted, leading to higher dysfluency rates at word, phrase, and sentence boundaries compared to within-word syllabic transitions (Bloodstein and Ratner, 2008; Buhr and Zebrowski, 2009). Neuroimaging studies indicate that atypical function in the cortico-basal ganglia circuits, including both the direct and indirect pathways, may contribute to speech disruptions (Alm, 2004; Giraud *et al*., 2008), and recent work highlights the HDP as crucial for inhibitory control and preparatory motor regulation in speech (Nambu, 2011; Neef *et al*., 2015). The HDP conveys rapid signals from frontal inferior cortical regions to the STN, enabling stopping and adjustment of ongoing motor commands, which supports sensorimotor integration and accurate coordination of speech movements (Whillier *et al*., 2018; Usler, 2022). Dysfunction in these circuits may impair the ability to reset or prepare the motor cortex at motor planning boundaries (e.g., word boundaries), providing a possible explanation for why stuttering often emerges at transitions between hierarchical speech units.

Although the focus of this study was on the segmental level of speech production, prosodic information can also define word boundaries. In this study, the syllables marking a word boundary also mark a difference in prosody. Other studies implicate the right IFG in the perceptual categorization of emotionally-valenced prosodic information (Sammler *et al*., 2015; Zhang *et al*., 2026). The right STG is also implicated in prosody and modulated by its emotional salience (Sammler *et al*., 2015), however, right STG condition-differences did not emerge in this study. One possibility is that the right IFG is part of a dorsal stream (Hickok and Poeppel, 2007; Ausilio, Craighero and Fadiga, 2012; Correia, Jansma and Bonte, 2015) associated with sensorimotor transformations, which can include information about prosody and may therefore inherently contribute to decisions about word boundaries. Nevertheless, the right IFG emerged in the comparison between the 2+4 and 3+3 conditions, where stress pattern was held constant. Hence, prosody differences cannot fully account for the right IFG activation differences found between conditions, here. The results are more likely consistent with a role for the right IFG in a basal-ganglia cortical loop that supports segmental sequences forming larger units, namely word units.

Several possible limitations should be considered. First, the sample size (N=19), although typical for EEG studies may limit sensitivity to smaller effects between condition 2+4 and 3+3, especially given the need to correct for multiple time-frequency bins tested per IC cluster. Second, within-word transitions were 17 milliseconds shorter between syllables 2 and 3, and 29 milliseconds shorter between syllables 3 and 4. These timing differences may also contribute to the ERSP estimates. Third, paced speech constrains generalization of the findings to more natural and prosodically richer speech production; but strict pacing and pseudo-words enhance control and timing, allowing a more controlled analysis of EEG data in epochs less likely contaminated by speech responses (i.e., speech -related EMG).. Finally, while the most parsimonious interpretation of the results in the right IFG cluster is related to the cortico-basal ganglia circuit underlying the HDP, we cannot fully rule out other factors confirming this as its sole role. Scalp EEG has limited coverage for subcortical regions (i.e., reduced signal-to-noise-ratio from deep brain sources) and thus does not provide a direct measure. In the future, simultaneous extracranial and intracranial EEG recordings that more directly target subcortical regions (e.g., from the STN in PD patients undergoing deep-brain stimulation - DBS) can provide additional insights into the interpretation of the results. Altogether, the behavioural and neural data indicate that hierarchical structure shapes both the timing and pre-speech cortical dynamics of speech production. The right-lateralised IFG oscillatory signature preceding the target syllable suggests predictive control mechanisms tuned to whether the upcoming syllable completes or starts a word, complementing accounts of hierarchical temporal scaffolding in speech perception and production (Ghitza, Giraud and Poeppel, 2013; Ding *et al*., 2015; Piai *et al*., 2015).

